# HCR-Proxy resolves site-specific proximal RNA proteomes at subcompartmental nanoscale resolution

**DOI:** 10.1101/2025.05.30.656369

**Authors:** Anja Trupej, Valter Bergant, Jona Novljan, Martin Dodel, Tajda Klobučar, Maksimiljan Adamek, Flora Lee, Karen Yap, Eugene Makeyev, Boštjan Kokot, Luka Čehovin Zajc, Andreas Pichlmair, Iztok Urbančič, Faraz Mardakheh, Miha Modic

## Abstract

The spatial organization of RNA condensates is fundamental for understanding of basic cellular functions, but may also provide pivotal insights into diseases. One of the major challenges to understanding the role of condensates is the lack of technologies to map condensate-scale protein architecture at subcompartmental or nanoscale resolution. To address this, we introduce HCR-Proxy, a proximity labelling technique that couples Hybridization Chain Reaction (HCR)-based signal amplification with *in situ* proximity biotinylation (Proxy), enabling proteomic profiling of RNA-proximal proteomes at subcompartmental resolution. We benchmarked HCR-Proxy using nascent pre-rRNA targets to investigate the distinct proteomic signatures of the nucleolar subcompartments and to uncover a spatial logic of protein partitioning shaped by RNA sequence. Our results demonstrate HCR-Proxy’s ability to provide spatially-resolved maps of RNA interactomes within the nucleolus, offering new insights into the molecular organisation and compartmentalisation of condensates. This subcompartment-specific nucleolar proteome profiling enabled integration with deep learning frameworks, which effectively confirmed a sequence-encoded basis for protein partitioning across nested condensate subcompartments, characterised by antagonistic gradients in charge, length, and RNA-binding domains. HCR-Proxy thus provides a scalable platform for spatially resolved RNA interactome discovery, bridging transcript localisation with proteomic context in native cellular environments.

**GRAPHICAL ABSTRACT:** 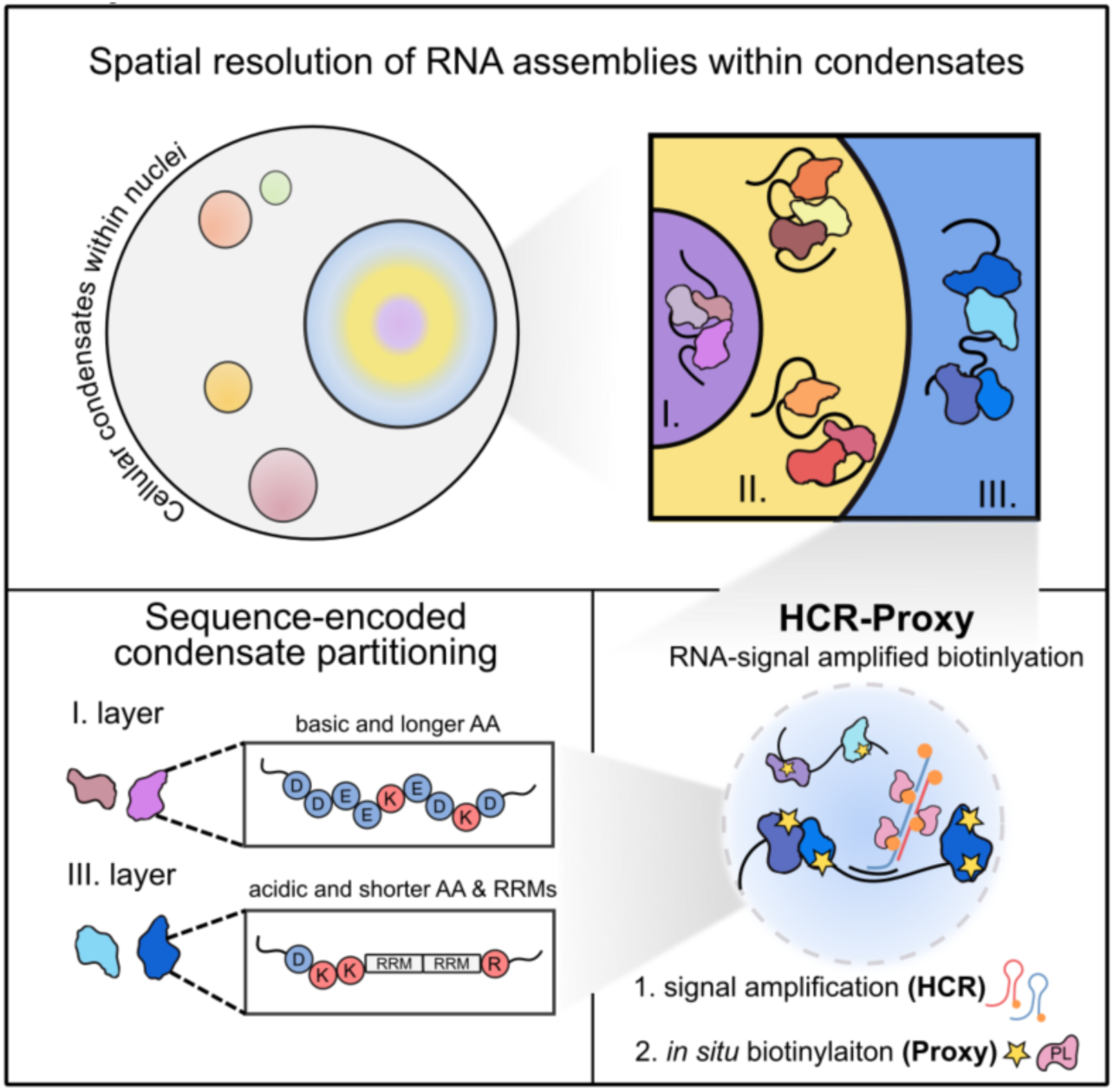

**Bullet points:**

- HCR-Proxy enables the first nanoscale-resolution mapping of RNA-proximal proteomes *in situ*.
- HCR-Proxy establishes a broadly applicable modular platform for spatially resolved RNA– interactomics.
- Subcompartmental proteomes are resolved across nucleolar subdomains by targeting specific pre-rRNA regions.
- Deep learning confirms a sequence-encoded logic of protein partitioning within condensate subcompartments.

## INTRODUCTION

The cellular milieu is organized into a myriad of membraneless compartments or biomolecular condensates, many of which are formed through multivalency-driven phase separation^1^. RNA molecules often serve as a nucleation seed, providing an architectural blueprint for the recruitment of RNA-binding proteins (RBPs)^2^ and ultimately condensate’s formation. Many condensates are heterogeneous entities, displaying various degrees of internal subcompartmentalization. Nucleoli^3^, Cajal bodies^4^, paraspeckles^5^ and nuclear speckles^6^ are amongst the best known examples of macro-assemblies with information workflow organized in nested-like structures^4^. Therefore, characterising molecular assemblies on subcompartment-wide scale is of vital importance for understanding the flux of biomolecules governed by the spatial organisation and most importantly its contribution to the cell’s homeostasis.

High-resolution microscopy using immunofluorescence is widely employed to study condensate composition^7–9^, while RNA-protein interaction mapping has relied on affinity-based approaches^10–12^, and newer RNA-targeted degradation methods^13,14^. Although these techniques identify direct interactors, they lack spatial context information crucial for compartment-level analysis. Proximity labelling (PL) overcomes this caveat by utilising an engineered biotinylating enzymes (e.g., APEX2^15,16^, HRP^17^, TurboID^18^), colocalising with the molecule of interest. Upon addition of a biotin substrate enzymes catalyse short-lived reactive species which are covalently deposited on biomolecules located in enzymes’ close proximity. Subsequently, biotinylated molecules are enriched by streptavidin pulldown and identified by mass spectrometry or nucleic acid sequencing^15^. Despite a broad repertoire of *in vivo* PL methods^19–23^, such approaches require genetic manipulations of the cells and exhibit low signal-to-noise ratio and experimental reproducibility. To overcome these limitations, *in situ* PL approaches^24–26^ have been developed that use labelled antisense oligos to biotinylate targeted RNA neighbourhoods without genetic editing. Despite bypassing the need for genetic manipulations, challenges remain in labelling specificity, spatial resolution and inefficient biotinylation, thereby preventing the characterisation of proximal interactome at subcompartmental scale. Thus, there is a pressing need to develop a biotinylation method with enhanced labelling and spatial resolution capacity to map spatially-confined RNA interactomes at the subcompartmental resolution.

To address this need, we have developed HCR-Proxy, an improved proximity labelling approach to identify site-specific RNA proteomes with subcompartment spatial resolution. HCR-Proxy is a hybrid method composed of a signal amplification (HCR) and an *in situ* biotinylation step (Proxy). Unlike current *in situ* PL methods, where biotinylation efficiency depends on the number of labelled probes recognised by the PL enzymes, HCR-Proxy uses metastable hairpins to amplify the biotinylation signal. This results in probe-independent RNA labelling that is suitable for studying proximal proteomes of challenging RNA molecules with limited sequence space for antisense probe design. To improve the resolution of the labelling, we concomitantly applied automated STED microscopy and profilled various molecular crowding conditions to technically refine the method. We benchmarked HCR-Proxy against different nuclear RNA targets of different abundances and demonstrated its flexibility and superior biotinylation efficiency. As a proof of concept, we applied HCR-Proxy to target specific nucleolar biogenesis-driven rRNA transcripts, demonstrating the method’s ability to specifically enrich distinct RNA-proximal proteomes within a nested condensate. Furthermore, the integration of deep learning models showed that basic protein features such as sequence length, composition and net charge can classify proteins located in different RNA-scaffolded neighbourhoods. Overall, this demonstrates that HCR-Proxy is a powerful tool with unprecedented spatial resolution for deciphering higher-order interactomes within spatially restricted RNA compartments. Thereby HCR-Proxy expands the toolbox for spatial proteomics by unifying RNA programmability with enzymatic labelling and biophysical modulation.

## RESULTS

### HCR-Proxy: Modular approach for targeted *in situ* proximity labelling through amplified RNA recognition

Several RNA *in situ* hybridisation techniques have been developed for visualisation of subcellular localization of individual RNA transcripts^27,28^. Among them, the *in situ* hybridisation chain reaction, or HCR, offers an unparalleled specificity, sensitivity and high signal-to-noise ratio^29,30^. Unlike traditional *in situ* hybridisation approaches, where signal intensity depends on the number of directly labelled probes hybridised to the target, HCR approach requires programmable oligonucleotide probes to hybridise in pairs to adjacent regions of the selected RNA. Specifically, bound probe pairs trigger the signal amplification by the self-assembling chain reaction of fluorophore-labelled hairpins. To date, this method has been applied for fluorescence *in situ* hybridisation (FISH)^29,31,32^.

Building on the strengths of HCR-FISH, we sought to extend the advantages of HCR signal amplification for a proximity labelling approach to precisely identify biomolecules interacting with or in close proximity to a specific RNA target. Therefore, we developed HCR-Proxy, a hybrid *in situ* proximity labelling workflow composed of signal amplification (**HCR**)^27,30^ and biotinylation step (**Proxy**) (Fig. 1A).

**Figure 1:**
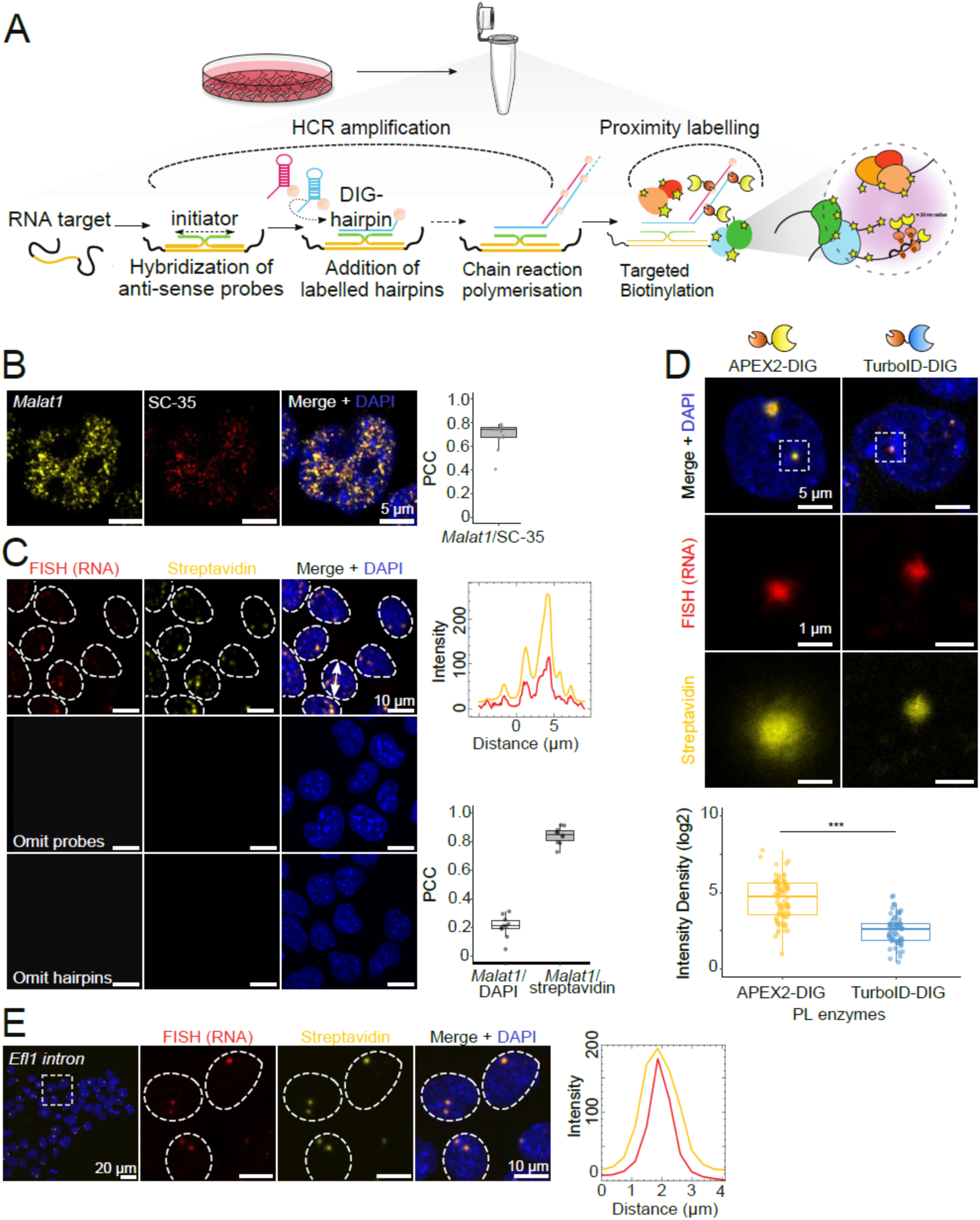
HCR-Proxy design and implementation. A) Schematic overview of HCR-Proxy. The main part of the in situ workflow is streamlined into tubes, where chemically fixed cells are hybridized with pairs of anti-sense probes towards target RNA. Only the complete probe pair alignment enables signal amplification (HCR) via polymerised DIG-labelled metastable hairpins and therefore recruitment of PL enzyme to catalyse in situ proximity biotinylation (Proxy). B) HCR-FISH / IF micrographs of non-coding RNA Malat1 (yellow, DyLight594-conjugated anti-DIG) in mESCs colocalising with nuclear speckle marker SC-35 (red, SC-35 antibody), merged with DAPI staining (blue). The colocalisation of the signal was calculated using Pearson’s correlation coefficient (PCC) from 13 images (dots represent each cell). C) HCR-Proxy FISH micrographs of Malat1 in mESCs supported with signal intensity profiles of RNA FISH (red, DyLight594-conjugated anti-DIG) and HCR-Proxy (yellow, Alexa647-conjugated streptavidin) signals. Cells nuclei were visualized with DAPI staining blue). The colocalisation of the signal was calculated using Pearson’s correlation coefficient (PCC) from 10 images (dots represent each cell). D) HCR-Proxy FISH micrographs of Efl1 intronic transcript (red, DyLight594-conjugated anti-DIG) and its proximal vicinity labelled with two different PL enzymes, APEX2 or TurboID (yellow, Alexa647-conjugated streptavidin), merged with DAPI staining (blue). Difference in intensity density was calculated with two-sided Student’s t-test (p < 0.001 = ***, p < 0.01 = **, p < 0.05 = *). E) HCR-Proxy FISH micrographs of Efl1 intronic transcript with signal intensity profiles of RNA FISH (red, DyLight594-conjugated anti-DIG) and HCR-Proxy (yellow, Alexa647-conjugated streptavidin) signals, merged with DAPI staining (blue).

First, we developed metastable digoxigenin-labelled hairpins (DIG hairpins) that specifically bind only the fully assembled initiator sequence to trigger the growth of a tethered digoxigenin amplification polymer (Fig. 1A). All non-specifically bound probes and hairpins stay kinetically trapped and therefore do not trigger amplification. The amplified number of DIG molecules at the target RNA serves as a binding pad for a recombinant PL enzyme, triggering proximity-labelling of adjacent biomolecules *in situ* while the unbound enzyme is washed off. To preserve biomolecular interactions we chemically fix cells with cell permeable and reversible cross-linker DSP^33,34^, an amine-group specific alternative to paraformaldehyde, which can distort interactions and thus change the condensate’s composition^35^. Upon permeabilisation, cells are transferred into tubes, where subsequent HCR-Proxy steps (Methods) are streamlined, making the protocol more user-friendly and cost-efficient. Finally, biotinylated proteins are enriched by magnetic streptavidin beads under denaturing conditions followed by identification with mass spectrometry (HCR-Proxy MS) (Fig. 1A).

To validate the HCR-Proxy methodology we targeted *Malat1* (Table S1), a hallmark non-coding RNA of nuclear speckles^36^, in mouse embryonic stem cells (mESCs). As expected, the HCR-FISH signal colocalised with the well-known nuclear speckle marker, SC35 (Fig. 1B)^7^. To assess the method’s biotinylation efficiency, we further performed proximity labelling and stained for fluorescently-labelled target RNA and surrounding biotinylated biomolecules. Pearson correlation analysis revealed strong spatial colocalisation with a high degree of signal overlap (Fig. 1C). To evaluate specificity of the method, we also conducted control experiments by omitting either the hairpin amplifiers or probe pairs. Both conditions resulted in the absence of FISH and streptavidin signals, demonstrating the requirement for complementarity effect between probes and amplifiers to achieve biotinylation (Fig. 1C). This demonstrates that HCR-Proxy can accurately proximity label RNA in its native cellular environment.

To evaluate the enhanced biotinylation efficiency achieved by signal amplification, we designed two sets of probes targeting the intronic region of *Efl1* mRNA (Table S1), localised to two nuclear puncta. First, we aimed to identify the PL enzyme that would provide the strongest and most specific biotinylation on lowly abundant RNA sites. To achieve this, we purified recombinant enzymes APEX2^25^ and TurboID^18^, fused to an optimised binder selective for digoxigenin (DIG binding domain)^37^ (Fig. S1). We observed an approximately 5.5-fold higher biotinylation efficiency with APEX2-DIG, yielding high biotinylation signal-to-noise ratio while maintaining specific colocalisation with the target RNA (Fig. 1D). These results demonstrate that HCR-Proxy methodology can be used in conjunction with a versatile range PL enzyme and indicates that HCR-Proxy efficiently biotinylates even proximity of lowly-abundant nascent RNAs using only two sets of probes (Fig. 1E).

### Enhancing spatial specificity through biophysical tuning of labelling conditions

Next, we implemented automated content-aware STED super-resolution microscopy^38^ to measure the biotinylation diameter *in situ* and to screen for conditions to improve sensitivity and resolution of proximity labelling reaction. Fully automated STED image acquisition platform with data-adaptive imaging, similar to implementation reported in this preprint^39^ (see Methods for details), provided an efficient capture of high-resolution biotinylation signal distributions, minimizing operator-induced variability and enhancing data reliability and reproducibility (Fig. 2A). Super resolution imaging revealed that biotinylation staining extended beyond the boundaries of *Efl1* RNA localisation, spreading within a micrometre range (Fig. 2B), consistent with previous reports of other PL approaches^25^. Surprisingly, rather than intensity peak overlapping with the RNA FISH signal, we detected a reduced biotinylation signal in the condensate core compared to its periphery, resulting in a "halo labelling effect" and reduced labelling specificity of cellular microenvironments (Fig. 2B).

**Figure 2:**
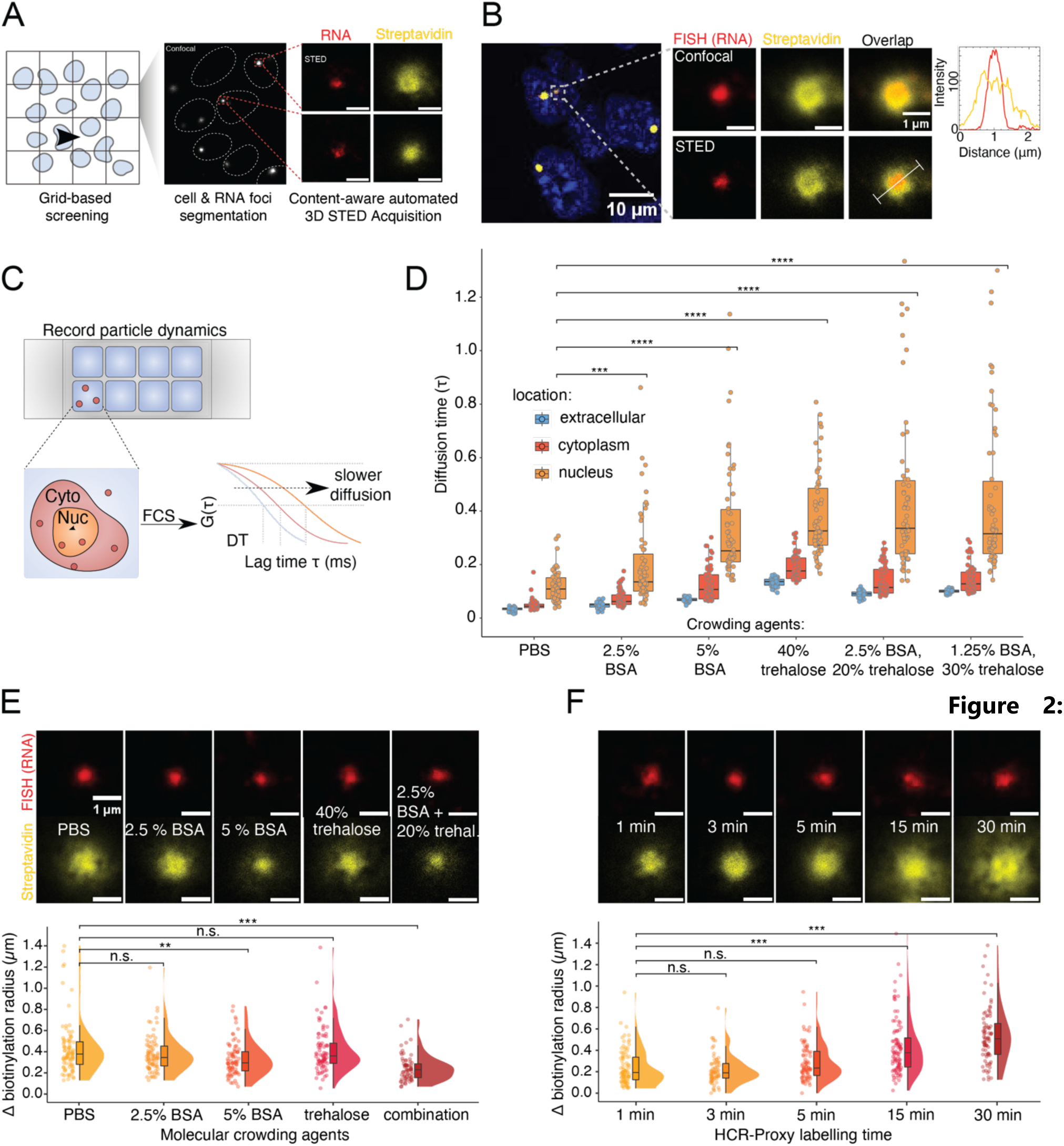
Spatially-restricted enhanced proximity labelling workflow limits the diffusion of activated biotin radicals. A) Schematic workflow of STED image acquisition platform. In each coarsely sampled confocal image of the tiled panorama scan, nuclei and RNA foci were automatically identified for subsequent super-resolution STED imaging. B) STED microscopy reveals nonspecific biotinylation. STED micrograph and intensity plot of the biotinylation signal (yellow, Alexa647-conjugated streptavidin) beyond RNA FISH boundary (red, DyLight594-conjugated anti-DIG). Reduced intensity of the biotinylation signal in the condensate’s core is barely detected with confocal microscopy. C) Schematic workflow of the fluorescence correlation spectroscopy (FCS) experiment to infer the particle diffusion dynamics (red dots present tracer molecule Alexa 647) sampled in cells’ nucleus, cytoplasm, and extracellular compartment. Autocorrelation curves shifted towards longer lag times indicate slower diffusion. Characteristic diffusion times (DT) shown in panel D were obtained by fitting experimental autocorrelation curves with a 3D diffusion model. D) Quantification of diffusion time for Alexa647 fluorophore from FCS experiment within different cellular compartments for different conditions. Average number of measurements per condition, depicted as dots: extracellular = 30, cytoplasm = 60, nucleus = 60 (two-sided Student’s t-test; p < 0.001 = ***, p < 0.01 = **, p < 0.05 = *). E) Effect of molecular crowding agents on the biotinylation specificity. Top: HCR-Proxy FISH micrographs against Efl1 intronic transcript, where proximity labelling was performed with PBS, 2.5 % BSA, 5 % BSA, 40 % trehalose and Combination of BSA and trehalose (2.5 % BSA & 20 % trehalose). Bottom: Quantification of the difference in radius size between biotinylation (yellow, Alexa647-conjugated streptavidin) and RNA FISH signal (red, DyLight594-conjugated anti-DIG). Approximately 100 images per condition were acquired, depicted as dots (two-sided Student’s t-test; p < 0.001 = ***, p < 0.01 = **, p < 0.05 = *). F) Effect of labelling time extension on the biotinylation specificity. Top: HCR-Proxy FISH micrographs of Efl1 transcript, where proximity labelling was performed under five different labelling times: 1, 3, 5, 15 and 30 minutes, respectively. For this experiment labelling solution Combination 1 was utilized. Bottom: Quantification of the difference (delta) in radius size between biotinylation (yellow, Alexa647-conjugated streptavidin) and RNA FISH signal (red, DyLight594-conjugated anti-DIG). Approximately 100 images per condition were acquired, depicted as dots (two-sided Student’s t-test; p < 0.001 = ***, p < 0.01 = **, p < 0.05 = *).

To overcome the challenge of nonspecific biotinylation, we next aimed to spatially constrain biotin radicals by utilising inert crowding agents that either increase the buffer viscosity (PEG^40^, various sugars^41^) or alternatively elevate the local protein concentration (BSA^41,42^). To assess whether these crowding agents reduce the diffusion of activated biotin radicals, we performed fluorescence correlation spectroscopy (FCS), which measures molecular diffusion dynamics based on intensity fluctuations as molecules diffuse through a defined volume of light^43^. Fixed and permeabilised cells as per HCR-Proxy protocol were stained with DAPI and different solutions of crowding agents were applied. PBS served as a reference labelling solution. To approximate biotin radicals behaviour, we used Alexa647 fluorophore and recorded the mobility of molecules within nuclei, cytoplasm and extracellular compartment. Fitting autocorrelation functions with a 3D diffusion model, we obtained the diffusion times (i.e. average time for molecules to diffuse through the focal volume) for each condition (Fig. 2C), which revealed a significant effect of crowding agents on diffusion time in all three compartments. As expected^41,42^, diffusion was slower within nuclei compared to other two compartments, regardless of labelling solution (Fig. 2D). These findings show that molecular crowding plays a crucial role in modulating diffusion dynamics and as such biotinylation signal. The strongest effect was observed with a combination of BSA and trehalose (Fig. 2D), leading to a significant decrease in diffusion rate. These results demonstrate that controlled modulation of diffusion dynamics can be achieved through tailored crowding conditions, providing a useful strategy for enhancing labelling specificity in PL experiments.

To further validate our findings, we performed an HCR-Proxy experiment using the same combination of labelling solutions. Using our automated STED screening platform, we assessed the effect of molecular crowding agents on biotinylation specificity by comparing the radius of RNA FISH and biotinylation signal (Fig. 2E). The most efficient biotin radical’s confinement was observed with a synergistic combination of BSA and trehalose at their lower concentrations. In addition to reducing labelling radius, crowding agents also abolished the “halo labelling effect”. We propose that the decrease in radical mobility not only spatially constrained radicals to the site of their production but also prevented the inactivation of PL enzyme.

Next, we investigated whether crowding agents could compensate for extension of labelling time without compromising specificity, thereby increasing biotinylation sensitivity and consequently higher downstream yield. To determine optimal labelling time, we performed HCR-Proxy FISH experiment at 5 different time points and measured the biotinylation radius surrounding the targeted RNA. We were able to extend the standard 1 min labelling time of HCR-Proxy PL to 3 or 5 minutes without any noticeable spread in biotinylation radius. However, exceeding the labelling time to 15 or 30 minutes led to a significant increase in biotinylation radius and reappearance of the “halo labelling effect” (Fig. 2F). These results indicate that fine-tuning the biotinylation efficiency by a combination of crowding agents and labelling time extension enhances the biotinylation yield without compromising specificity, thereby limiting the *in situ* proximity labelling reaction to individual microenvironments of biomolecular condensates.

Upon technically refining labelling and crowding conditions, we next aimed to benchmark HCR-Proxy to characterise the proximal proteomes of individual RNA microenvironments. For this we performed HCR-Proxy LC-MS (liquid chromatography coupled to mass spectrometry) in mESCs with probes targeting precursor ribosomal RNA (pre-rRNA) and ncRNAs *Malat1*, *Norad*, along with intronic *Efl1* sequence (Table S1, Fig. S2A, Fig. 1B). This allowed us to quantify levels of 1130 proteins with varying abundances across different HCR-Proxy targets. Consistent with the distinct cellular localisation of the targets, Uniform Manifold Approximation and Projection Analysis for dimensionality reduction (UMAP) showed a clear separation of samples between nuclear and nucleolar targets (Fig. S2B). We considered proteins as *bona fide* nucleolar proteins if they were significantly enriched over remaining nuclear targets (Student’s t test-based log2 fold-change > 1, FDR-adjusted p-value < 0.05) (Fig. S2C), thereby obtaining 602 nucleolar proteins, including numerous well-established pre-rRNA interactors (Tables S2, S3). Consistent with the data, GO term annotation of the pre-rRNA proximal proteome also showed an enrichment for ribosomal proteins (Fig. S2D). With this, we have shown that HCR-Proxy facilitates target-dependent enrichment of protein interactors of RNA and thus is capable of profiling RNA-scaffolded compartments.

### Resolving nested RNA microenvironments with HCR-Proxy

With the unprecedented specificity (Fig. 2), we postulated that HCR-Proxy for the first time paves the way towards spatially-resolved proximity labelling of RNP condensates. To demonstrate the versatility of HCR-Proxy in deciphering the interactome within individual condensates, we aimed to delineate the multiphased composition of nucleoli, which consists of (at least) three subcompartments^44^. For this we targeted different regions of the nascent rRNA transcript, (i) upstream or (ii) downstream of A’ processing and (iii) spanning ITS2 region (Table S1), localised in the fibrillar centre (FC), the dense fibrillar component (DFC) and the granular component (GC), respectively (Fig. 3A). Using HCR-FISH staining, we observed mutually exclusive signals for all three targeting sites (RNA baits) (Fig. 3B), confirming their distinct spatial distribution within the nucleolus.

**Figure 3:**
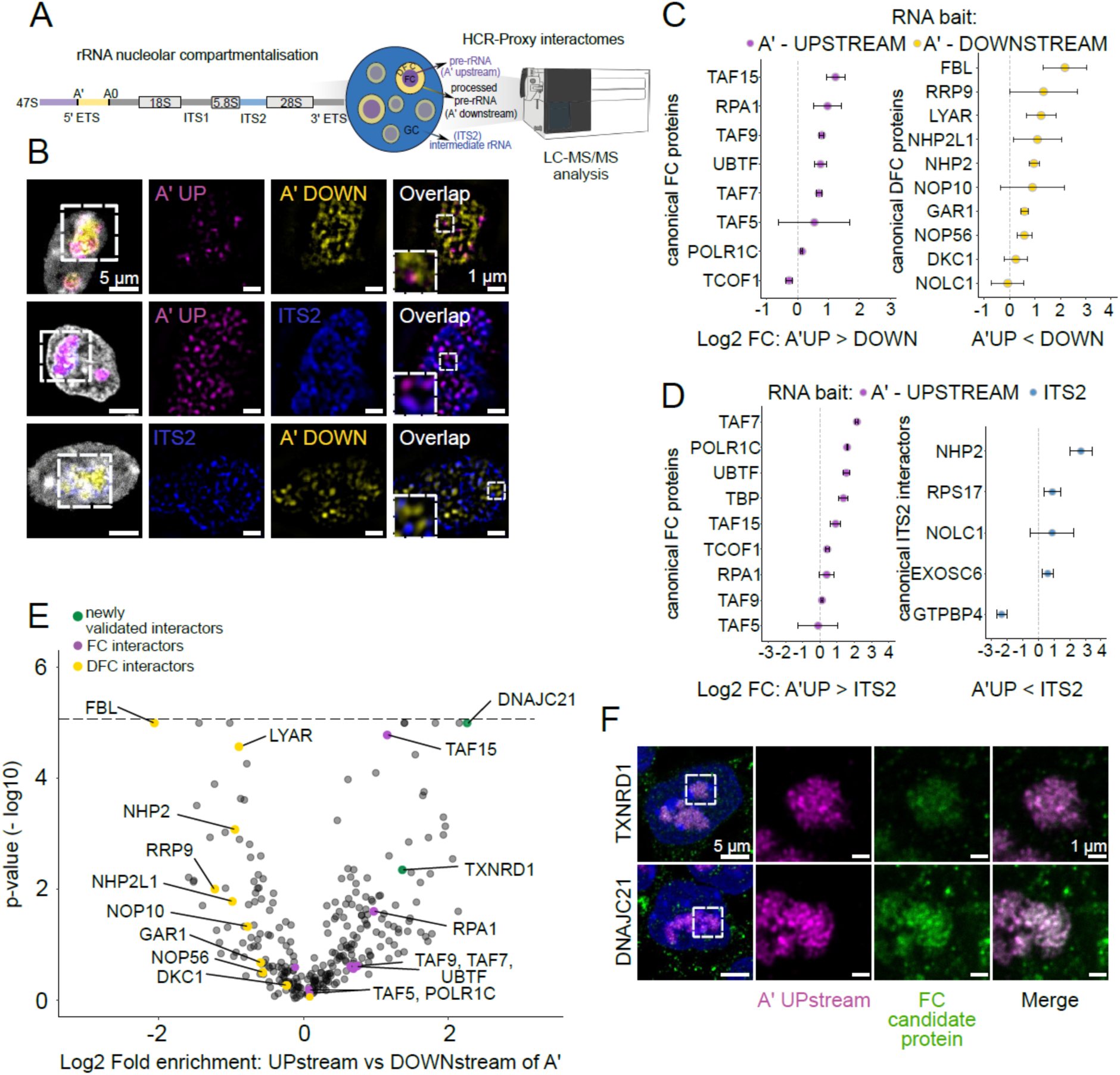
Spatially resolved rRNA-proximal proteomes within subnucleolar subcompartments. A) Schematics of probes designed for HCR-Proxy MS to spatially resolve pre-rRNA proteomes within multiphased nucleoli. B) HCR-FISH deconvoluted STED micrographs of probes targeting distinct regions of the nascent rRNA transcript (magenta - A’ upstream, yellow - A’ downstream, blue - ITS2). C) and D) Dot plots displaying enriched subnucleolar HCR-Proxy components, demonstrating ability of HCR-Proxy MS to spatially resolve multiphased nucleolar interactomes between (C) A’ upstream and A’ downstream and (D) A’ upstream and TS2 region, respectively. Error bars present standard deviations. E) Volcano plot of enriched interactors specific for A’ upstream region (magenta), its newly validated interactors (green) and A’ downstream region (yellow). The cut-off for identification of bona fide interactors was set on Lasso regularization-based log2 fold-change > 1 and associated -log10(p-value) > 1. For visualisation purposes, -log10(p) values (y-axis) were thresholded at 5 (dashed line). F) HCR-FISH / IF STED micrographs of newly validated A’ upstream interactors. RNA FISH (magenta - A’ upstream) and newly validated nucleolar proteins localised to FC (green - DNAJC21, TXNRD1).

At the same time, each of them also colocalised with its known subnucleolar marker (FC (POLR1A), DFC (FBL), GC (NPM1)) (Fig. S3A). In agreement with the nucleolar localisation of all three pre-rRNA targets, HCR-Proxy MS identified distinct subnucleolar proximal proteomes (Table S4). When comparing the HCR-Proxy interactomes for nascent pre-rRNA regions upstream (UP) and downstream of A’ processing site (DOWN), we detected prominent components for rRNA transcription and established FC markers enriched in the UP region. The later enrichment includes numerous transcription factors responsible for forming Pol I pre-initiation complex like UBTF^9,45,46^, TAF15, TAF9, TAF7 and TAF5^47^, alongside with POLR1C component of the RNA Pol I complex (Fig. 3C)^9^ and RPA1 protein that is involved in rRNA transcription regulation^48^. In contrast, numerous proteins involved in early stages of pre-rRNA processing, including FBL^9,49^, its interaction partner GAR1^49^, LYAR and various components of the box C/D and H/ACA snoRNP complexes such as NHP2, NOP56 and NOP10^9,46^ were enriched in the HCR-Proxy interactome specific for the DOWN region (Fig. 3C). We were also able to distinguish the recently identified ITS2 interactors^13,50^ when compared to the UP region, where we recapitulated some of the known components responsible for its processing such as RPS17, NHP2, NOLC1 and EXOSC6 (Fig. 3D).

Encouraged by the recovery of many known interactors specific to each pre-rRNA region, we next sought to investigate whether HCR-Proxy could reveal previously unreported subnucleolar components. To this end we selected two candidate proteins enriched in HCR-Proxy for UP region bait: cytosolic heat shock protein DNAJC21 and thioredoxin reductase TXNRD1, none of which had been previously characterised within the nucleolus (Fig. 3E). Both interactors exhibited strong colocalisation with RNA FISH signal, with distinct high-intensity peaks overlapping the UP bait region (Fig. 3F). These results combined indicate that HCR-Proxy not only distinguishes interactomes from spatially restricted RNA assemblies within the same condensate, but also offers a powerful strategy for uncovering novel subnucleolar components.

### Sequence-encoded logic of condensate partitioning based on HCR-Proxy interactomes

We next postulated that with the expansion of the pool of identified proteins for each compartment, we can now fully leverage deep learning (DL) models to further refine our understanding of the sequence-encoded determinants that govern protein partitioning across different nucleolar subcompartments. To achieve optimal performance and interpretability, we benchmarked three deep learning frameworks under stratified three-fold cross-validation (Fig. 4A) in classifying the A′-upstream or A′-downstream region-specific interactome. The first model we implemented is a convolutional neural network (Sequence-CNN) trained directly on full-length amino acid sequences, designed to recognize local patterns and sequence motifs predictive of the target classification. The second model (ESM2-MLP) leverages embeddings derived from the transformer-based protein language model ESM-2^51^, aiming to capture global context and higher-order interactions among amino acids across the entire sequence. Finally, the third model (Feature-LogReg) relies on 320 diverse manually curated features (Fig. 4A), with sequence-derived properties calculated using localCIDER^52^ and NARDINI^53^, and annotated characteristics obtained from UniProt^54^. A greedy-forward-selection distilled this large feature set to identify the 14 most predictive descriptors of compartment identity, used in the final model.

**Figure 4:**
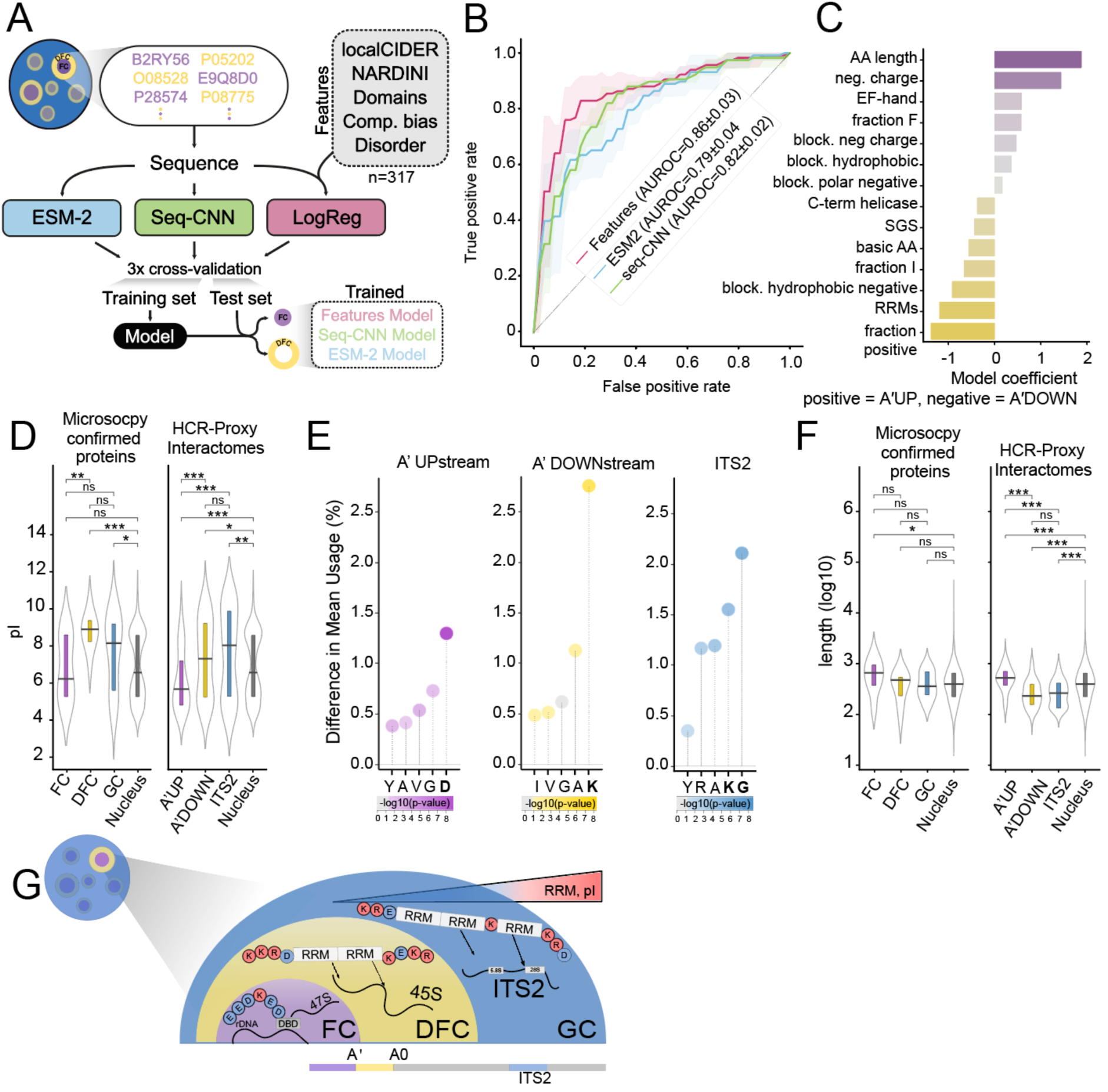
Protein sequence code distinguishes HCR-Proxy obtained nucleolar subcompartmental interactomes. A) Schematic overview of the deep learning models trained to classify A′-upstream (FC-associated) and A′-downstream (DFC-associated) interactomes. Protein sequences were classified using either ESM-2 embeddings, a sequence-based CNN model (Seq-CNN), and a logistic regression model based on 320 protein features (see Methods). Models were evaluated using 3-fold cross-validation. B) Receiver operating characteristic (ROC) curves showing the classification performance of each model, with mean area under the curve (AUROC) and standard deviation across cross-validation folds. C) Coefficient analysis of the logistic regression model identifying the top predictive features of A′UP vs A′DOWN localization. Positively weighted features (purple) are enriched in A′UP proteins; negatively weighted features (yellow) are enriched in A′DOWN proteins. D) Comparison of the isoelectric point (pI) between nucleolar proteins confirmed by microscopy and identified by HCR-Proxy MS. Number of proteins FC = 15, DFC = 13, GC = 28 *^9^*, A’ UP = 118, A’ DOWN = 84, ITS2 =75, GO-termed nuclear = 15585 (Mann–Whitney U-test; p < 0.001 = ***, p < 0.01 = **, p < 0.05 = *). E) Comparison of amino acid usage in A′ UP, A′ DOWN and ITS2 proximal proteomes relative to the nuclear background. The five residues with the largest differences in mean usage are highlighted. Dot colour reflects statistical significance (Welch’s t-test: p < 0.05 = *, p < 0.01 = **, p < 0.001 = ***). F) Comparison of amino acid length between nucleolar proteins confirmed by microscopy and identified by HCR-Proxy MS. Number of proteins FC = 15, DFC = 13, GC = 28 *^9^*, A’ UP = 118, A’ DOWN = 84, ITS2 =75, GO-termed nuclear = 6023 (Mann–Whitney U-test; p < 0.001 = ***, p < 0.01 = **, p < 0.05 = *). G) Graphical summary of protein features across subnucleolar compartments.

Surprisingly, the Feature-LogReg classifier attained an average AUROC of 0.86, outperforming the protein language model based ESM2-MLP classifier (AUROC = 0.79) and the Sequence-CNN (AUROC = 0.82) (Fig. 4B). Coefficient analysis revealed that HCR-Proxy-enriched proteins in the A′-upstream interactome (associated with FC) are characterised by greater protein length and elevated acidic residue count. In contrast, HCR-Proxy-enriched proteins in the A′-downstream interactome (associated with DFC) exhibit a higher fraction of basic residues and a pronounced enrichment of RNA-recognition motifs, consistent with the presumed role of DFC proteins in mediating pre-rRNA processing (Fig. 4C).

Machine-learning–driven feature selection revealed an antagonistic pattern of net charge, RNA-binding modules, and protein length between the FC and DFC proteomes. To validate these predictions, we first calculated isoelectric points (pI) for all HCR-Proxy targets along with all remaining GO-annotated nuclear proteins. Consistent with the feature model’s identification of increased acidity, the FC cohort exhibited a significantly lower median pI than both the DFC and GC groups, and versus the nuclear reference (Fig. 4D). Per-residue analysis confirmed that Asp (D) frequencies were markedly elevated in FC proteins, whereas Lys (K) dominated the DFC and GC proteomes (Fig. 4E, S3B). These sequence-derived features of HCR-Proxy interactomes are strongly supported by experimental dissection of protein sequences of canonical nucleolar proteins harbouring D/E tracts and K-blocks^46^. For instance, the prominence of blocky acidic residues (Fig. S3B, C) among FC-enriched proteins echoes their identification of acidic D/E tracts as a determinant of FC localisation and lower local pH. Similarly, the authors identified the characteristic K-blocks regions within DFC scaffolds, which we validated to be enriched (Welch’s t-test, p < 0.001) in HCR-Proxy A’ downstream interactome (associated with DFC) (Fig. S3C).

Next, we analysed the protein domains characteristics for nucleolar subcompartments. Annotation of RNA-recognition motifs showed a marked increase in RRM domain content from FC to DFC and GC (Fig. S3D), which aligns with the presumed role of DFC/GC proteins in mediating pre-rRNA processing and RNA binding domains being excluded from FC^46^. Finally, we compared protein lengths and found that DFC and GC proteins were on average roughly half as long as FC or other nuclear proteins (Fig. 4F), indicating that DFC proteins tend to be shorter, basic, with higher RNA-binding capacity and rich in lysine-rich blocks important for RNA interaction and processing^55^.

In conclusion, only the rich, compartment-specific proteomic data enabled by HCR-Proxy allowed us to deconvolve the molecular code underlying nucleolar organisation. This data-driven insight not only identified distinct sequence features driving spatial partitioning, but also lends strong support to the pH gradient model^46^ wherein longer, acidic FC proteins scaffold to the fibrillar core, whereas shorter, basic, RRM-enriched proteins preferentially localise to the surrounding dense fibrillar and granular components to orchestrate dynamic rRNA processing (Fig. 4G).

## DISCUSSION

Biomolecular condensates have emerged as critical hubs of cellular organisation, with compartmentalisation often orchestrated through RNA-mediated multivalent interactions. While considerable progress has been made in cataloguing the global composition of various condensates^56^, the spatial architecture of condensate patterning or proteomic mapping within nested condensates - such as the nucleolus - have been constrained by limitations in both resolution and sensitivity of RNA-centric proteomics approaches.

To address this challenge, we developed HCR-Proxy, a proximity labelling method that leverages hybridisation chain reaction (HCR) for both signal amplification and spatial confinement of biotinylation, thereby enabling nanoscale characterisation of RNA-proximal proteomes *in situ*. By designing DIG-labelled HCR hairpins as localised docking platforms for recombinant DIG-binding enzymes, HCR-Proxy introduces a modular and plug-and-play architecture: any recombinant enzyme fused to a DIG-binding domain (such as APEX2 or TurboID in this work) can be readily recruited to the RNA target site. This modularity allows broad adaptability across different labelling chemistries and also circumvents the constraints of conventional genetically encoded PL systems, allowing for seamless application in primary cells and complex systems where genome editing may not be feasible.

One of further innovations in our workflow is the confinement of labelling reactions to nanometre-scale environments through diffusion tuning by employing macromolecular crowding agents. When combined with automated content-aware 3D STED imaging to identify optimal buffer viscosity, this approach enabled unbiased, high-resolution mapping of labelling radii and supported the development of HCR-Proxy as a biophysically tuneable system for high-specificity labelling within nanoscale RNA-defined niches. Despite the diffusion confinement of labelling reactions, efficiency of biotinylation with HCR-Proxy does not require a high number of hybridisation probes, but is rather the result of signal amplification achieved by DIG-labelled hairpins at the site of the targeted RNA. This enabled us to use as little as two probe sets to achieve biotinylation of proximal proteomes, even for intronic RNA targets (Fig. 1E).

Using this framework, we demonstrate that HCR-Proxy can resolve sub-nucleolar microenvironments within a single condensate. As a hallmark multilayered condensate we used nucleolus and targeted distinct regions of pre-rRNA nascent transcript, each representing one of the three nucleolar sublayers. This enables molecular dissection of spatial proteome sublayers within a single condensate, revealing not only known markers but also uncovering novel nucleolar proteins with subnucleolar precision. In this way, we expanded the repertoire of identified proteins for each subcompartment and integrated deep learning classifiers trained on amino acid sequences, biophysical features and domain architectures to provide a functional link between sequence-encoded features and localisation to distinct rRNA-defined subdomains. Consistent with recent findings that subcellular localisation is governed by distributed “protein codes”^57^, our findings also reinforce the biophysical model of pH gradients within the nucleolus^46^, where acidic IDRs containing D/E tracts scaffold the FC, creating a microenvironment distinct from the lysine-rich, RRM-rich proteins of the DFC and GC. Next, our deep learning analysis also revealed that simple, interpretable features such as protein length, net charge, and domain content can outperform transformer-derived embeddings in classifying subnucleolar localisation. This suggests that phase behaviour and spatial partitioning are deeply encoded in primary protein sequence and this predictive logic extends prior findings from engineered systems to endogenous interactomes.

## DATA AVAILABILITY

Newly produced data was deposited to ProteomeXchange with identifier PXD064165. The code and notebooks to analyse the data and to produce the figures in this work are available at: https://github.com/ModicLab/HCR_proxy. The code used for HCR-Proxy image acquisition is deposited at Zenodo (10.5281/zenodo.15489876) and so are acquired microscopy images: (10.5281/zenodo.15479057).

## AUTHOR CONTRIBUTIONS

Conceptualisation: A.T., M.M,.; Project Administration: A.T., M.M., Investigation: A.T., V.B., J.N., M.D., T.K., M.A.,B.K.; Formal Analysis: A.T., V.B., J.N.,, B.K., I.U., F.M., M.M.; Writing - Original Draft: A.T., J.N, M.M.; Writing - Review Editing: A.T., V.B., J.N., E.M., F.L., F.M., I.U., M.M.; Methodology: A.T., V.B., J.N., M.M.; Data Curation: A.T., V.B., J.N., M.D.; Software: J.N., B.K., L.C.Z.; Supervision: E.M., A.P., I.U., F.M., M.M.; Resources: K.Y., E.M., A.P., I.U., F.M., M.M.; Funding Acquisition: A.P., F.M., M.M

## ACKNOWLEDGEMENTS

We are grateful to Leo Kurian for useful comments on the manuscript. This research was supported by Carl Zeiss Foundation (CZS Center SynGen), the Slovenian Research Agency (J4-50145, J4-60070, J7-2596, N1-0240, J4-60082, P1-0060), Janko Jamnik PhD Fellowships awarded to TK and MA, Helmholtz Program “Natural, Artificial, and Cognitive Information Processing”, King’s Prize of King’s College London (Wellcome & Mellows Charitable Settlement) and Johanna Quandt Young Academy fellowship hosted by Vascular Research Center, Faculty of medicine, Goethe University Frankfurt. HPC VEGA is financed through HPC RIVR consortium (www.hpc-rivr.si) and EuroHPC JU (eurohpc-ju.europa.eu). AP is funded by the Bavarian State Ministry of Science and Arts (Prevention of Pandemic-infection-associated Pathology Munich - P3M; BayVFP 2024-2027), the German Research Foundation (DFG) (TRR237 (A07), TRR179 (TP11), TRR353 (B04)) and the Danish National Research Foundation (DNRF 164; CiViA).

## MATERIAL AND METHODS

### Cell culture

Low-passage, wildtype mouse pluripotent stem cells IDG3.2 PSCs (129S8/B6 background^58^) and endogenously tagged Polr1a-gfp mESCs were cultured in a humidified incubator at 37 °C, 5 % CO2, feeder-free, on cell culture dishes (TPP) coated with 0.1% Gelatin (Millipore, cat# ES-006-B). Cells were maintained in N2B27 medium composed of 1:1 Neurobasal (Thermo Fisher Scientific, cat #21103049) and DMEM-F12 (Thermo Fisher Scientific, cat #11320074) medium, containing N2 (Thermo Fisher Scientific, cat# 17502001) and B27 (Thermo Fisher Scientific, cat# 17504001) supplements, 12 ng/mL LIF (Qkine, cat #104278), with additional use of small molecule inhibitors: for condition commonly named 2iLIF, 1 mM MEK inhibitor PD0325901 (Axon Medchem, cat# 1408) and 3 mM GSK3 inhibitor CHIR99021 (Sigma, cat# SML1046). Cells were passaged every 2-3 days using Accutase (Sigma, cat# A6964).

### Generation of CRISPR/Cas9 genome engineering mESC lines

For the generation of Polr1a-GFP specific gRNAs targeting upstream of the stop codon (gRNA sequence AGACAATGCTGCTATCTTAG) were cloned into a modified version of the SpCas9-T2A-GFP/gRNA plasmid (px458172, Addgene plasmid #48138), where we fused a truncated form of human Geminin (hGem) to SpCas9 in order to increase homology-directed repair efficiency generating SpCas9-hGem-T2A-GFP/gRNA. To generate Polr1a-GFP targeting donors, dsDNA gene fragments were synthesized and cloned into a vector carrying ∼400 bp homology arms (IDT). For targeting, wild-type ESCs were transfected with a 4:1 ratio of donor oligo and SpCas9-hGem-T2A-GFP/gRNA construct. Positively transfected cells were isolated based on GFP expression using fluorescence-activated cell sorting (FACS) and plated at clonal density in ESC media 2 days after transfection. After 5–6 days, single colonies were picked and plated on 96-well plates. These plates were then duplicated 2 days later and individual clones were screened for the desired mutation by PCR followed. Cell lysis in 96-well plates, PCR on lysates, and restriction digests were performed. The presence of the desired Polr1a-GFP insertions in putative clones was confirmed by Sanger sequencing.

### Recombinant proximity labelling enzymes

#### MOLECULAR CLONING

Plasmid pML433^59^ for bacterial production of fusion protein APEX2-DIG with N-terminal T7 tag, flexible linker between APEX2 and DIG, and C-terminal TEV-cleavage site and hexahistidine tag was a kind gift from Dr Karen Yap, King’s College London, UK. In this plasmid, APEX2 is inserted into the plasmid via EcoRI and HindIII restriction sites, which were used to replace APEX2 with TurboID. The gene fragment for TurboID was taken from Addgene Plasmid #107177 and purchased from IDT with 5’ EcoRI and 3’ HindIII restriction sites. The TurboID gene fragment and pML433 with APEX-DIG were double digested with FastDigest EcoRI (cat# FD0274) and FastDigest HindIII (cat# FD0505) (Thermo Fisher Scientific). The restriction reactions were resolved on a 1 % AGE gel. pML433 without APEX2 and the digested TurboID gene fragment were isolated from gel with the use of Zymoclean Gel DNA Recovery Kit (Zymo Research) and further purified with the use of Zymoclean DNA Clean & Concentrator-25 Kit (Zymo Research). T4 DNA ligase (Thermo Fisher Scientific, cat# EL0011) was then used to ligate TurboID into pML433 in an overnight reaction performed at 16 °C. The ligation mixture was transformed into chemically competent *E. coli* cloning strain DH5ɑ via standard heat shock procedure. Bacteria were plated on a LB agar plate supplemented with 30 μg/mL kanamycin (LBK-agar). After overnight incubation at 37 °C, colonies were analysed via colony PCR with the use of DreamTaq PCR Master Mix (Thermo Fisher Scientific, cat# K1081) and T7-promoter and T7-terminator annealing oligonucleotides (purchased from IDT). Positive clones were transferred into liquid LB media supplemented with 30 μg/mL kanamycin (LBK) and grown overnight (37 °C and shaking at 150 rpm). The following day, plasmid DNA was isolated with the use of ZR Plasmid Miniprep - Classic Kit (Zymo Research, cat# D4015). Isolated DNA was sent for Sanger sequencing (Macrogen). The clone with the correct sequence and best A_260_/A_280_, A_260_/A_230_ ratios (as measured by NanoDrop One, Thermo Fisher Scientific) was chosen for subsequent work.

#### BACTERIAL CULTURE

Plasmid pML433 with construct T7-tag_APEX2_linker_DIG_hexahistidine-tag (APEX2-DIG) or T7-tag_TurboID_linker_DIG_hexahistidine-tag (TurboID-DIG) was transformed into chemically competent *E. coli* expression strain BL21(DE3) via standard heat shock procedure. Bacteria were plated on an LB agar plate supplemented with 30 μg/mL kanamycin. After overnight incubation at 37 °C, a single bacterial colony was transferred to 10 mL of LB medium containing 30 μg/mL kanamycin. The culture was incubated with shaking at 37 °C and 150 rpm for 8 hours. Then, 100 µL of starter culture was transferred to 50 mL of fresh LBK. The culture was left overnight with shaking at 37 °C and 150 rpm. Next day, 2.5 mL of overnight culture was added to 250 mL of TB media supplemented with 100 mM phosphate buffer (pH 7.0) and 30 μg/mL kanamycin (TBK). For the APEX2-DIG construct, the media was also supplemented with 0.1 mM FeSO_4_. The cultures were grown with shaking at 37 °C and 150 rpm to an OD_600_ of 1.0. The cultures were briefly cooled on ice, induced with 0.5 mM IPTG, and left overnight (∼18 h) with shaking at 20 °C and 150 rpm. Next day, cultures were centrifuged, supernatant discarded, and the resulting bacterial cell pellets were frozen at −80 °C until use.

#### PROTEIN PURIFICATION

Protein purification was done following a slightly modified protocol established by Yap and colleagues^59^. Briefly, bacterial cell pellet (5 g for APEX2 construct and 3 g for TurboID construct) was left to thaw on ice. Then, ice-cold lysis buffer (50 mM HEPES (pH 8.0), 200 mM NaCl, 10 mM MgCl_2_, 10 % glycerol, 2.5 mM βME, 1 mM PMSF, 0.1 % (V/V) triton X-100, 250 U benzonase, 0.5 mg/mL lysozyme) was added (pellet:buffer m/V ratio = 1:10) and the pellet was gently resuspended. The cells were lysed by sonication on an ice bath (Cole Parmer Ultrasonic Processor; 38 % amplitude, 1 sec pulse time, 2 sec pause, total pulse time 5 min). The lysed cells were centrifuged at 50,000 rcf for 30 min at 4 °C. The supernatant was filtered through a 0.22 µm filter. Protein purification was performed on an ӒKTA pure M25 chromatography system (Cytiva), which was kept in a fridge at 4 °C. The clarified lysate was loaded onto a HR 10/20 Ni-NTA Superflow (Qiagen) column, which was pre-equilibrated with IMAC buffer A (25 mM HEPES (pH 8.0), 200 mM NaCl, 5 % glycerol, 2.5 mM βME). The column was washed with stepwise increasing concentration of IMAC buffer B (25 mM HEPES (pH 8.0), 200 mM NaCl, 5 % glycerol, 2.5 mM βME, 500 mM imidazole): 2.5 % IMAC buffer B (10 CV) and 5 % IMAC buffer B (10 CV). Elution was carried out with 50 % IMAC buffer B. Protein peak from IMAC was loaded onto a HiLoad Superdex 200 PG 16/60 column (GE Healthcare), which was pre-equilibrated with SEC column buffer (25 mM HEPES (pH 8.0), 200 mM NaCl, 5 % glycerol, 1 mM DTT). Chosen fractions were analysed by SDS PAGE (mPAGE 4-12 % Bis-Tris precast gels, Merck Millipore, cat# MP41G10) and fractions containing high amounts of purified protein were pooled together. Protein concentration was estimated by NanoDrop One. APEX2-DIG construct was stored as is (concentration 2 mg/mL), while TurboID-DIG construct was concentrated with the use of a Pierce Protein Concentrators PES, 10K MWCO (Thermo Fisher Scientific), to a final concentration of 1 mg/mL. Proteins were aliquoted and stored at −80 °C.

Semiquantitative tests for APEX2-DIG peroxidase activity and its digoxigenin binding ability were performed by the protocol previously described ^25^. TurboID-DIG biotinylation activity was determined by HCR-Proxy FISH staining.

### DNA probes and DIG-labelled hairpins for proximity labelling

DNA oligonucleotide split probes (Table S1) complementary to precursor ribosomal RNA (A’ upstream), processed pre-rRNA or A’ downstream (A’ - A0), intermediate rRNA (ITS2), Malat1, Norad and Efl1 were designed by a custom software (ÖzpolatLab-HCR, 2021 - Github^32^). Sequences generated by the software were purchased from IDT, resuspended to 100 µM with nuclease-free water and mixed for each probe stock with the final 5 µM concentration. Digoxigenin and fluorophore-labelled HCR hairpins (Table S1) were custom designed by Molecular Instruments Inc. and stored in a light-tight container at - 20 °C.

### Immunofluorescence and HCR-FISH

Hybridisation-chain reaction FISH was performed according to the protocol^30^. WT IDG3.2 mESCs were plated on Geltrex™ hESC-Qualified (Thermo Fisher Scientific, cat# A1569601) coated 8-well glass-bottom IBIDI plate one day prior to fixation. Cells were fixed with 0.5 mg/mL dithiobis(succinimidyl propionate) (DSP; Thermo Fisher Scientific, cat# 22585) in 1 × PBS for 40 min at room temperature, washed three times with 1×PBS and 20 mM Tris-HCl, pH 7.5, 5 minutes each wash and permeabilized with 70 % ethanol overnight at 4 °C. Hybridisation and amplification steps were performed as described in the protocol. For immunofluorescence stainings, samples were after amplification thoroughly washed three times with 5x SSCT (5x SSC (Invitrogen, cat# 15557044), 0.1 % Tween-20), 5 minutes each wash, two times with 1x PBS and blocked with IF blocking buffer (0.3 % Triton-X-100 + 3 % BSA + 10 U/mL Rnasin (Promega, cat# N2615)) for 1 h at room temperature. To visualize RNA labelled with digoxigenin we used DyLight® 594 anti-DIG antibody (goat, Vectorlabs, cat# DI-7594-.5, 1:100 dilution). To validate colocalisation with nuclear condensates and confirm localisation of candidate proteins we used the following antibodies against: SC-35 (mouse, Santa Cruz Biotechnology, cat#sc-53518, 1:200 dilution), NPM1 (mouse, ThermoFisher Scientific, cat# 32-5200, 1:100 dilution), FBL (rabbit, Abcam, cat#ab5821, 1:200 dilution), TXNRD1 (rabbit, Proteintech, cat# 11117-1-AP, 1:200 dilution), DNAJC21 (rabbit, Proteintech, cat# 23411-1-AP, 1:100 dilution) and GFP (rabbit, ThermoFisher Scientific, cat# A-21311, 1:100 dilution). Samples with antibodies were incubated with 0.8 % BSA in 4x SSC (Blocking buffer) supplemented with 10 U/mL Rnasin inhibitors overnight at 4 °C. Next day, samples were washed with 4x SSC, 4x SSC and 0.1 % Triton X-100 and 4xSSC, 10 minutes each wash, and incubated with secondary antibodies, anti-mouse Alexa Fluor 647 conjugated (donkey, Invitrogen, cat# A-31571, 1:400 dilution) or anti-rabbit Alexa Fluor 647 conjugated (donkey, Abcam, cat# ab150075, 1:400 dilution) for 1 h at room temperature. Afterwards, cells were washed with 4x SSC, 4x SSC and 0.1 % Triton X-100 and 4xSSC, 10 minutes each wash, and cell nuclei were stained with DAPI (ThermoFisher, cat# 62248, 1:3000 dilution) in 1x PBS for 8 minutes at room temperature. Samples were mounted with 250 μL of Fluoromount G (Thermo Fisher Scientific, cat# 00-4958-02), sealed with parafilm and saved at 4 °C until imaging.

### HCR-Proxy

For HCR-Proxy MS experiments approximately 5 million cells per replicate were used in four replicates per each RNA target (bait) for subcellular HCR-Proxy targeted interactomes.

Cells grown in 10-cm dishes (HCR-Proxy MS) or 8-well glass-bottom IBIDI coverslips (HCR-Proxy FISH / IF staining) (IBIDI) (90 % confluency) were washed one time with 1x PBS, aspirated and fixed with 0.5 mg/mL DSP in 1 × PBS for 40 min at room temperature. Samples were afterwards washed three times with 1×PBS and 20 mM Tris-HCl, pH 7.5, 5 minutes each wash and permeabilized with 70 % ethanol overnight at 4 °C. Next day cells were rehydrated with two washes of 2x SSC, scraped into DNA LoBind tubes (Eppendorf, cat# 30108051) with 2xSSC supplemented with 1 % BSA (pluriSelect Life Science UG, cat# 60-00020-11) and 0.5x phosSTOP (Sigma Aldrich, cat# 4906837001) and centrifuged for 5 min, with 3400 rpm at room temperature to remove the supernatant. From this point on all steps were carried out within DNA LoBind tubes, rotated on the rotor and after each incubation or wash centrifuged for 5 min, 3400 rpm at room temperature to remove any residue. HCR hybridization and amplification steps were performed by following the HCR-FISH protocol^30^. Additionally, to prevent RNA degradation and other enzymatic activity 1x phosSTOP and 10 U/mL Rnasin inhibitors were added to pre-hybridisation and pre-amplification steps, meanwhile 1x phosSTOP and 25 U/mL Rnasin inhibitor were supplemented to tubes for overnight incubations such as probe hybridisation and HCR (hairpin) amplification.

After HCR amplification, cells were centrifuged and washed 3x times with 5x SSCT for 5 minutes at room temperature to remove the unbound hairpins. Cells were blocked with the Blocking buffer (0.8 % BSA in 4x SSC) and supplemented with 50 U/mL Rnasin inhibitor for 30 min at room temperature. Afterwards cells were incubated with 2.7 µg/mL TurboID-DIG or APEX2-DIG PL enzyme in the Blocking buffer for 1h at room temperature on the rotor. To remove any unbound PL enzyme, cells were thoroughly washed by 1x time 4x SSC, 2x times with 4x SSC + 0.1 % Triton and 2x times with 4x SSC for 10 min each. Next, cells were transferred to 2 mL DNA LoBind tubes (Eppendorf, cat# 30108078) and incubated in 0.5 mL of Labelling solution (2.5 % BSA & 20 % trehalose (AMSBIO EUROPE B.V., cat# AMS.TS1M-100)) for 5 min on the rotor. Proximity labelling was then performed by the addition of an equal volume of ice-cold Labelling solution supplemented with 1 mM biotin-phenol and 0.2 mM hydrogen peroxide (Merck, cat# H1009-5ML) and gentle rotation on ice for a defined amount of time. To stop biotinylation reaction, cells were quenched three times with 1 mL of ice-cold Quencher solution (10 mM sodium ascorbate (Sigma, cat# SI-A4034-100G) and 5 mM Trolox (Sigma, cat# AL-238813-1G) in 1xPBS) and thoroughly shaken. Last wash was performed in 1.5 mL DNA LoBind tubes with an additional final centrifugation step to remove any remaining Quencher solution. Samples labelled in tubes were then analysed by immunoblotting and mass spectrometry, meanwhile coverslips were used for HCR-Proxy FISH/IF imaging.

To evaluate the effect of crowding agents on biotinylation radius, five different Labelling solutions were prepared: 2.5 % BSA, 5 % BSA, 40 % trehalose, 2.5 % BSA & 20 % trehalose and 1.25 % BSA & 30 % trehalose. All solutions were prepared in 1x PBS, meanwhile 1x PBS served as a biotinylation positive control.

### Isolation of biotinylated proteins

Biotinylated cells were lysed within tubes with 0.3 mL of High SDS RIPA lysis buffer (150mM NaCl, 1mM EDTA, pH 8.0, 50mM Tris-HCl, pH 8.0, 1 % NP40, 0.5% sodium deoxycholate, 0.5% SDS) supplemented with 1x phosSTOP + 1x cOmplete EDTA-free protease inhibitor (Sigma, cat# SRO-5056489001) + 10 mM sodium ascorbate + 5 mM Trolox +50 mM DTT (ZellBio, cat# DTT25), transferred into Bioruptor tubes (Diagenode, cat# C30010016) and incubated on ice for 15 - 20 min while resuspended on a shaker. Samples were sonicated using a Bioruptor Pico system (Diagenode) with a built-in cooling system (Bioruptor® Cooler), for 15 cycles of 30 s ON / 30 s OFF at the low frequency setting. Lysates were then transferred to new DNA LoBind tubes, topped to 1 mL with High SDS RIPA lysis buffer supplemented with 1x phosSTOP + 1x cOmplete EDTA-free protease inhibitor + 10 mM sodium ascorbate + 5 mM Trolox +50 mM DTT and reverse crosslinked on thermoblock at 37 °C for 60 min with 1500 rpm shaking. Each sample was additionally supplemented with 2 µL of Turbo DNase (2 U/μL, Invitrogen, cat# AM2238) and 1 µL of Benzonase (≥ 250 U/µL, Merck, cat# 101654). For thorough DNA denaturation and cell homogenisation we pressed the cell lysis through the 27G syringe (8-10 x times), followed by boiling the lysate for 5 min at 95 °C and centrifuged at 15,000 x g for 10 min at 4 °C. Clear supernatant was transferred to a new tube and stored at - 70 °C until needed.

Before Streptavidin pulldown, samples were diluted with additional 0.5 mL of High-SDS RIPA lysis buffer supplemented with 1x phosSTOP and pre-cleared with Dynabeads Protein G (Invitrogen, cat# 10004D). For pre-clearing, 15 µL of Dynabeads per sample were washed three times with iCLIP lysis buffer (50 mM Tris-HCl, pH 7.4, 100 mM NaCl, 1% Igepal CA-630, 0.1% SDS, 0.5% sodium deoxycholate), then resuspended with reverse-crosslinked lysates and incubated for 45 minutes at 4⁰C with rotation. Cleared lysates were collected using a magnetic rack and then incubated overnight at 4 ⁰C with rotation with 40 µL of Pierce Streptavidin magnetic beads per sample (Thermo Fisher Scientific, cat# 88817), pre-washed three times with iCLIP lysis buffer. The following day, beads were pelleted and subjected to sequential washes: three times with an iCLIP lysis buffer supplemented with 0.5 % SDS; two times with High salt washing buffer (50 mM Tris-HCl pH 7.4, 1 M NaCl, 1 mM EDTA, 1 % Igepal CA-630, 0.1 % SDS, 0.5 % sodium deoxycholate); one time with 2 M urea in 10 mM Tris-HCl, pH 8.0 for 1 minute; and two times with iCLIP lysis buffer (0.5 % SDS). For the final wash, beads were transferred to a new tube and washed five times with Wash buffer (50 mM Tris-HCl pH 7.4, 5 % glycerol, 1.5 mM MgCl2, 100 mM NaCl) to remove any residual SDS. After the last wash, beads were transferred to a new tube and resuspended in 37 µL of the Wash buffer. Out of this, 5 µL of them were taken for immunoblotting and the rest of the beads were put on the magnetic rack to remove any residual buffer and stored on - 70 ⁰C till the mass spectrometry sample preparation.

### MS sample preparation and data acquisition

Protein-loaded beads underwent bead digestion, where they were resuspended in 50 μL of 8 M urea (Merck, cat# U1250) and 50 mM ammonium bicarbonate (Merck, cat# A6141) (ABC buffer). Reduction was performed by adding DTT to a final concentration of 10 mM followed by incubation at 25°C with agitation of 1200 rpm. After 30 min, samples were alkylated by adding iodoacetamide (VWR, cat# 786–228) to a final concentration of 55 mM and incubated in the dark at 25 °C with agitation of 1200 rpm for another 30 min. Next, a 50 mM ABC buffer was added to each sample for a final volume of 200 μL and a urea concentration of 2 M. Overnight tryptic digestion was then performed by adding 0.5 μg of Trypsin/sample (Merck, cat# T6567) and an incubation at 25 °C, 1200 rpm. The next day, the supernatant was separated from the beads using a magnetic tube rack. After mixing the supernatant with 4 % acetonitrile (J.T.Baker, cat# 9012), 1 %TFA (Thermo Fisher, cat# 85183) (STOP4 buffer) at a ratio of 1:1, samples were desalted using the Stage Tip procedure ^60^ and recovered in 0.1% TFA, 0.5% acetic acid (Honeywell, cat# 33209), 2% acetonitrile (A* buffer) for MS analysis.

LC-MS analysis was performed on a Q Exactive-plus Orbitrap mass spectrometer coupled with a nanoflow ultimate 3000 RSL nano HPLC platform (Thermo Scientific), using a 50 cm × 75 μm RSLC C18 column (Thermo Fisher) and a 123 min gradient of 3% to 35% of Buffer B (0.1 % FA in acetonitrile) against Buffer A (0.1 % FA in LC–MS gradient water). A total of 6 out of 7 microliters were injected, and mass spectrometry was carried out as previously described ^62^.

### Immunobloting

Five µL of protein-loaded beads in the Wash buffer were mixed 1:1 with 50 mM DTT and 4x NuPAGE LDS Sample Buffer (Invitrogen, cat# NP0007) and boiled for 10 min at 95 °C to elute proteins. The samples were centrifuged, placed on a magnetic rack, and eluates were collected and analysed by SDS-Page using NuPAGE® Novex 4-12% Bis-Tris gels (Thermo Fisher Scientific, cat# NP0322BOX), followed by an electrotransfer to nitrocellulose membranes using the Trans-Blot Turbo Transfer System (Bio-Rad) according to manufacturer’s instructions. The membranes were blocked in 3 % BSA and 0.1 % Tween-20 for 1 h on the rotor at room temperature, then incubated for another 90 minutes with IRDye® 800CW Streptavidin (Li-COR, cat# 926-32230, 1:1000 dilution). Membranes were washed three times with PBS and 0.1 % Tween-20, with each wash lasting 10 minutes, and biotinylated proteins were visualized with iBright™ FL1500 imaging system (Thermo Fisher Scientific).

### HCR-Proxy FISH

Proximity-labelled samples on 8-well IBIDI coverslips (prepared as described above) were washed twice for 5 minutes each with PBS and incubated overnight at 4 °C with DyLight® 594 antibody (Vectorlabs, cat# DI-7594-.5, 1:100 dilution) in Blocking buffer supplemented with 10 U/mL Rnasin inhibitor. The following day, samples were washed with 4x SSC, 4x SSC and 0.1 % Triton X-100 and 4xSSC, 10 minutes each wash, and incubated with Streptavidin STAR RED dye (Abberior, cat# STRED-0120-1MG, 1:1000 dilution) in a Blocking buffer supplemented with 10 U/mL Rnasin inhibitor at room temperature for 1 h. Afterwards, cells were washed with 4x SSC, 4x SSC and 0.1 % Triton X-100 and 4xSSC, 10 minutes each wash, and cell nuclei were stained with DAPI (1:3000 dilution) in 1x PBS for 8 minutes at room temperature. Samples were mounted with 250 μL of Fluoromount G, sealed with parafilm and saved at 4 °C till imaging. Images for Fig. 1C and 1E were acquired with Leica TCS SP5 inverted laser scanning microscope on a Leica DMI 6000 CS module equipped with an HCX Plane-Apochromat lambda blue 63× objective and a numerical aperture of 1.4 (Leica Microsystems). Rest of microscopic images were acquired with STED microscopy described below.

### Fluorescence super-resolution imaging (STED)

Images were acquired with a customised STED microscope (Abberior instruments) using a 60× water immersion objective. We excited fluorescently labelled RNA and Streptavidin dye (biotinylation site) with pulsed lasers at 561 and 640 nm, respectively, and DAPI stained nuclei with a CW 405 nm laser. We acquired the fluorescence intensity through a confocal pinhole set to 0.9 AU using avalanche photodiodes with 500-550 nm, 580-625 nm or 655-720 nm filters (Semrock).

Automated acquisition was driven by our bespoke Python scripts generated for this manuscript^63^, which ran over a grid of 10x10 positions for each sample in the 8-well slide. At each position, a large field-of-view image with coarse sampling was acquired in the confocal mode to identify positions with FISH signal within the nucleus (overlap with DAPI signal). For each identified nuclear condensate, an axial 3D STED cross-section image was acquired to determine the precise focus for the final 2D STED image (resolution improved laterally, xy). Typically, the laser powers were around 2 and 20 μW for the 640 and 561 nm laser, respectively, and 90–140 mW for the STED laser. The image size was 3–7 μm, pixel size was set to 30 nm, pixel dwell time 10 μs. Detection was gated (gate delay 250 ps, width 8 ns), and signal was accumulated over 2–10 line repetitions to achieve a good enough signal. For both cross-sections also, a confocal image was acquired for comparison.

In total, the automated acquisition allowed us to capture 1000 STED images of individual condensates (an average of 100 per sample) within 12 hours. For each condition, approximately 80 % of the images that met the image criteria (not recognized as an artefact) were selected for further analysis.

### Image analysis

Acquired images were analysed in Fiji/ImageJ^64^ using custom-made macros. Colocalisation analysis was performed per cell, where image contrast for both compared channels/measurements was enhanced (saturated = 0.35), except when using Z-stack images, followed by colocalization analysis with the BIOP JACoP plugin. To quantify biotinylation efficiency for enzyme modularity, masks for RNA were created, applied to raw images of Streptavidin dye and afterwards the intensity of overlapped area was measured. To quantify the effect of crowding agents and time extension on biotinylation radius, masks for RNA and Streptavidin signal were created, RNA signal was then subtracted from the biotinylation area and divided by 2. For image deconvolution we applied Diffraction PSF 3D plugin with adjusted parameters (Index of refraction, NA, wavelength, image size, slice spacing and number of slices), followed by usage of Parallel Spectral Deconvolution^65^ plugin with Generalized Tikhonov (reflexive) method and customized regularization parameters. All statistical analyses were performed in R program.

### Fluorescence correlation spectroscopy (FCS)

WT IDG3.2 mESCs were plated on Geltrex™ hESC-Qualified coated 8-well glass-bottom IBIDI plate one day prior to fixation. Cells were fixed with 0.5 mg/mL DSP in 1 × PBS for 40 min at room temperature, washed three times with 1×PBS and 20 mM Tris-HCl, pH 7.5, 5 minutes each wash and permeabilized with 70 % ethanol overnight at 4 °C. The next day, cells were rehydrated with two washes of 2x SSC, followed by three washes of 1x PBS, 5 minutes each wash, cell nuclei were stained with DAPI (1:3000 dilution) in 1x PBS for 8 minutes at room temperature and then rinsed twice with 1x PBS. Cells were left in different labelling solutions, sealed with parafilm and saved at 4 °C till the fluorescence correlation spectroscopy experiment. As a tracer molecule 80 nM Alexa 647 NHS was used. We acquired the FCS data with the above-described Abberior Instruments microscope steered with our automated python script, which moved across wells with different samples and determined regions for the FCS recordings within a nucleus (based on the DAPI signal), in the cytoplasm (i.e. in the immediate vicinity of the nucleus), and outside the cells. Within each chosen spot, 3 FCS measurements were taken, recording the fluorescence fluctuations for 30 s with sampling step 2 μs. Excitation power of the pulsed 640 nm laser was set to 5–10 μW at the sample plain. For each sample, at least 50 measurements for each location were performed; i.e. for the 6 wells, almost 1000 FCS recordings were autonomously acquired over the course of approx. 10 hours.

The time traces were autocorrelated and analysed using a 3D diffusion model with a triplet component with the open source FoCuS-scan software (https://github.com/dwaithe/FCS_scanning_correlator)^66^. The fitted diffusion transit times are plotted in Fig. 2C-D.

### MS data analysis

The MS raw files were searched using MaxQuant 1.6.3.3 against the mouse (*Mus musculus*) Proteom Fasta file extracted from UniProt (2023). Default MaxQuant settings were used with the exception of enabling LFQ, Match between runs and re-quantify options. MaxQuant output was analysed using R (version 4.3.0). Log2-transformed LFQ values were used for differential protein abundance analysis. Proteins only identified by site, reverse matches, potential contaminants, and protein quantified in less than 3 (Fig. S2B, C) or all (Fig. 3C, D, E) replicates of any condition were removed from the analysis. Missing values were imputed by sampling from normal distribution derived from valid measurements, which was down-shifted by 1.8 standard deviations and squeezed by a factor of 0.3.

Dataset consisting of targeting pre-rRNA and ncRNAs *Malat1*, *Norad*, along with intronic *Efl1* sequence, was analysed in the following manner: protein abundances from pre-rRNA samples were compared to a union of those from *Malat1*, *Norad* and *Elf1* using two-sided Student’s t-test. Thus, obtained p-values were FDR-corrected. The following criteria were used for considering proteins as significant nucleolar proteins: enrichment in pre-rRNA over the other targets > 1 (log2 fold change) and FDR-adjusted p < 0.05.

Dataset employing probes against A’ upstream, A’ downstream and ITS2 regions also included different labelling times and was analysed using linear modelling based on LASSO statistics. Only protein groups, quantified through at least 2 distinct peptides, were used for the analysis.

The following experiment design was used for differential protein abundance analysis: log2(LFQ) ∼ *labtime* + *labtime:bait*, where *labtime* refers to labelling time (either 1, 3 (only measured for A’ downstream), 5, or 15 minutes), and bait refers to HCR-Proxy targets (either A’ upstream, A’ downstream or ITS2). The following effects were thus estimated: the effect of time-labelling across all targets relative to 1-minute labelling; the labelling-time dependent enrichment of protein interactors for A’ downstream and ITS2 relative to A’ upstream. The estimation of LASSO model parameters was performed using R package glmnet ^67,68^ (version 4.0.2) with thresh = 1e-28, maxit = 1e7, and nfolds = 11. The exact model coefficients and lambda value at cross-validation minimum (lambda.min) were extracted and used for P-value estimation by fixed-lambda LASSO inference using the R package selectiveInference ^69^, version 1.2.5. Default parameters were used with the following modifications: tol.beta = 0.025, alpha = 0.1, tailarea_rtol = 0.1, tol.kkt = 0.1, and bits = 100. The bits parameter was increased to 300 or 500 if the convergence was not reached. The sigma was explicitly estimated using function estimateSigma from the same package. No multiple hypothesis P-value correction was performed since that is facilitated by the unbiased selection of optimum lambda - and thereby the model - for each analysed protein separately.

### GO term enrichment analysis

Enrichment of nucleolar-localized proteins was assessed against a background of all UniProt-annotated nuclear proteins using the STRING database (v12.0)^70^, considering only the overlap as target. The top 10 enriched Gene Ontology Cellular Component (GO:CC) terms, ranked by lowest false discovery rate (FDR), were selected and mapped onto the GO-basic ontology graph to construct a directed acyclic subgraph with their ancestral nodes. Non-enriched parent terms were pruned to streamline the visualisation, with node size scaled to the observed protein count and node colour graded by −log10(FDR).

### Machine learning

To identify distinguishing characteristics between HCR-Proxy defined A ‘DOWN- and A’ UP-class proteins, we developed three supervised learning frameworks based on distinct protein feature representations.

In the first approach, we extracted sequence embeddings from the 33rd hidden layer of the ESM 650M language model ^71^. These embeddings were averaged across residues to yield a fixed-length, 1280-dimensional vector for each protein. A fully connected neural network was then trained using these vectors as input, comprising two hidden layers with dimensions equal to 80 and 40, respectively (using ReLU activation), followed by a softmax output layer. Training was performed using the Adam optimizer (learning rate = 1 × 10⁻⁴) and categorical cross-entropy loss. Early stopping based on validation loss (patience = 25 epochs, maximum = 10,000 epochs) was applied.

In the second framework, full length amino acid sequences were one-hot encoded, padded to the longest protein and passed through a deep convolutional neural network. The architecture consisted of five convolutional blocks with 128 filters with kernel size = 5, each followed by max pooling of size 2. A global average pooling layer aggregated features across positions, which were then passed to a 64-unit dense layer with dropout before final softmax classification. Optimization and training strategy were identical to the dense model above.

The third model used 317 curated protein-level features computed using the localCIDER ^52^ and NARDINI ^53^ toolkits, together with annotated UniProt protein features ^54^. Single value sequence features derived from localCIDER quantified physical and chemical sequence properties including the relative abundance of each canonical amino acid, fraction of charged residues (FCR), net charge per residue (NCPR), isoelectric point, molecular weight, mean hydropathy, and others. In parallel, NARDINI computed z-scores for non-random clustering of specific residue classes based on 100,000 composition-matched sequence scramblings, including diagonal and off-diagonal pairwise blockiness metrics of eight residue classes (polar, hydrophobic, positively charged, negatively charged, aromatic, alanine, proline, glycine). Functional annotations from UniProt were also incorporated to quantify the percent of the protein covered by known domains, compositional bias regions, and predicted disordered segments. All features were z-normalised and used to train a logistic regression classifier with L2 regularization (maximum 1,000 iterations). To reduce model complexity while retaining predictive power, a greedy forward feature selection using beam search (beam width = 10, improvement AUROC score tolerance = 1 × 10⁻³) was applied to the full feature set (320). This procedure identified a 14-feature subset that maximized average cross-validated AUROC. The final selected features were: count of negatively charged residues, fraction of positively charged residues, clustering of hydrophobic and negative residues, percent of protein with RNA-recognition motif, fraction of isoleucine residues, sequence length, fraction of phenylalanine residues, percent of protein with helicase C-terminal domain, clustering of hydrophobic and alanine residues, fraction of basic residues, clustering of negative and alanine residues, percent of protein with EF-hand domain, clustering of polar and negative residues, and percent of protein with SGS domain.

Training of each of the three models was conducted using stratified 3-fold cross-validation. For deep learning models, the best of ten replicate training runs per fold was selected based on validation AUROC score; logistic regression was fit once per fold. With feature-based models being most predictive, importance of each feature was quantified by mean absolute model coefficient values across folds.

## SUPPLEMENTARY FIGURES LEGENDS

**Supplementary figure 1:**
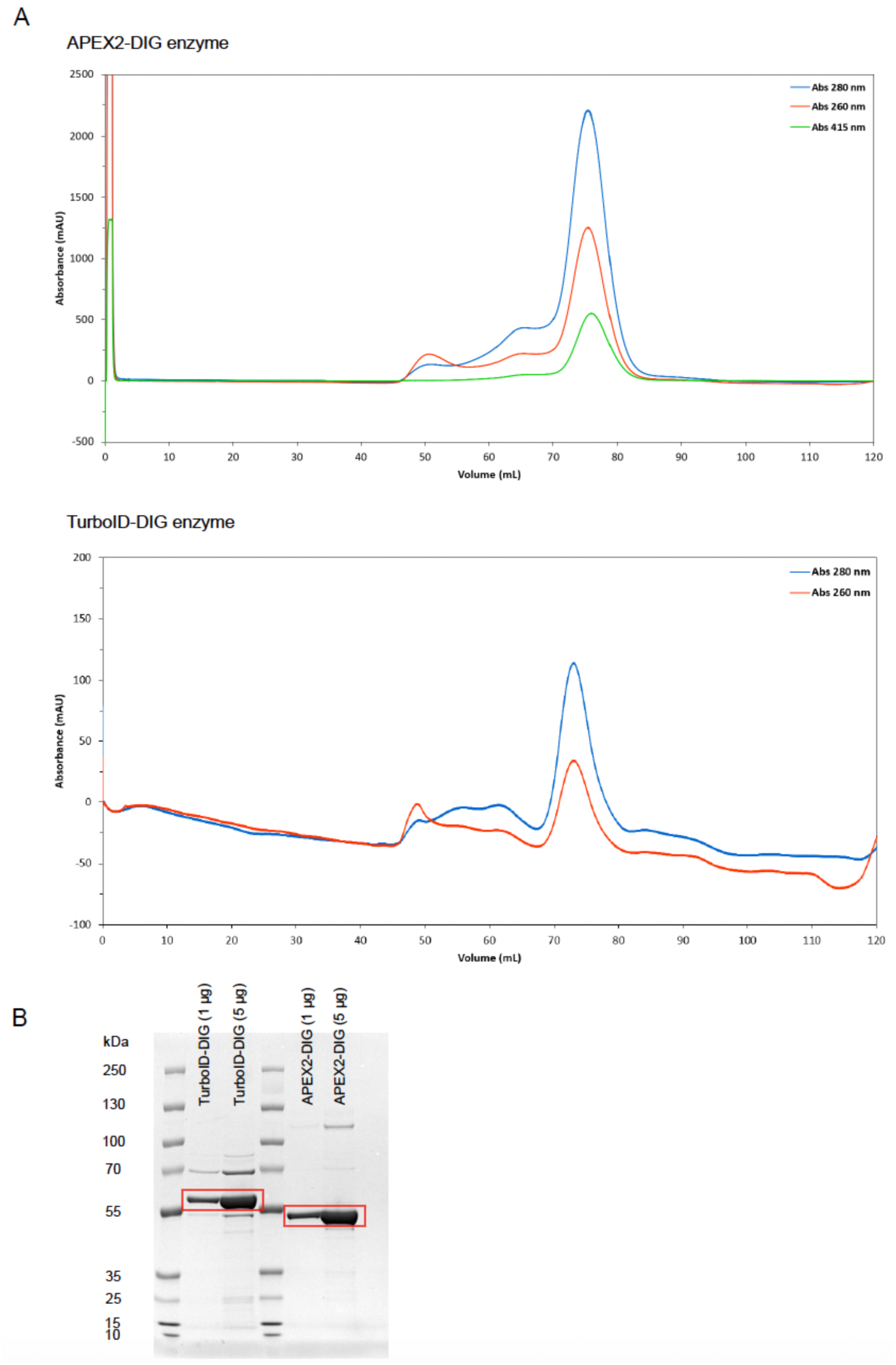
Isolation of PL enzymes TurboID-DIG and APEX2-DIG used for HCR-Proxy. A. Elution profiles from size exclusion chromatography for isolated PL enzymes. B. SDS-PAGE analysis of purified PL protein peaks eluted from a size exclusion chromatography column. Bands within red rectangles represent isolated enzymes.

**Supplementary figure 2:**
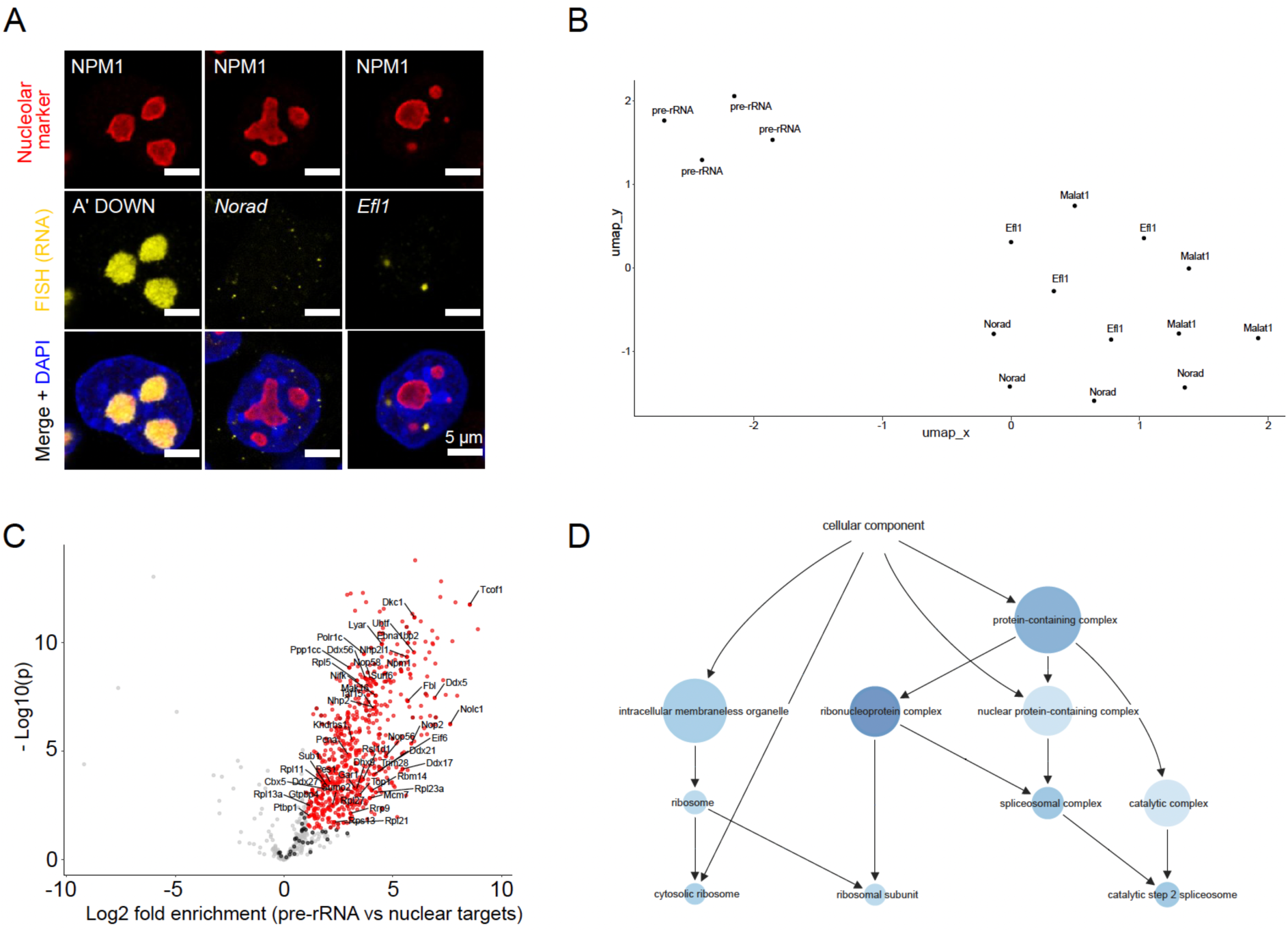
HCR-Proxy enables profiling of different RNA-scaffolded compartments within the cell. A. HCR-FISH micrographs of pre-rRNA (A’ downstream), Norad and intronic Efl1 sequence (yellow, DyLight594-conjugated anti-DIG) combined with immunofluorescence detection of nucleolar marker NPM1 (red, NPM1 antibody), merged with DAPI (blue). B. UMAP analysis of nucleolar and nuclear targets from benchmarked HCR-Proxy MS experiment. C. Volcano plot for the pre-rRNA-proximal proteome. Proteins marked in red present bona fide nucleolar proteins, meanwhile literature-known nucleolar proteins are labelled (two-sided Student’s t test-based log2 fold-change > 1, fdr- adjusted p-value < 0.05). D. GO term annotation of identified bona fide nucleolar protein interactors.

**Supplementary figure 3:**
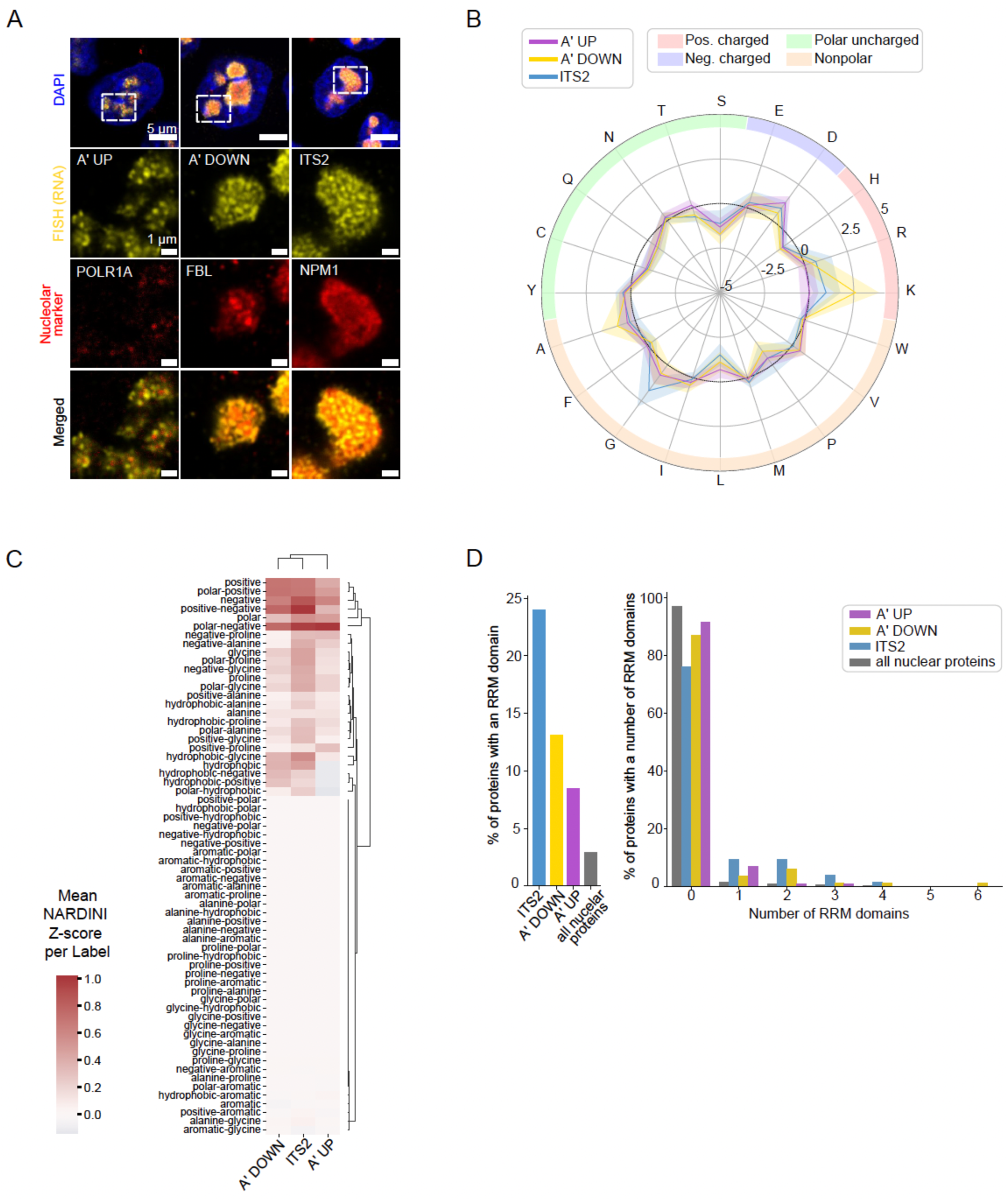
Primary sequence-derived features enable distinction between HCR-Proxy interactomes. A. HCR-FISH micrographs of RNA baits targeting sublayers of multiphased nucleolus (A’ upstream, A’ downstream and ITS2) (yellow, DyLight594-conjugated anti-DIG) combined with immunofluorescence detection of FC marker POLR1A, DFC marker FBL and GC marker NPM1 (red), merged with DAPI (blue). B. Radar plot showing the mean difference in amino acid usage in A′ UP, A′ DOWN, and ITS2 proximal proteomes relative to the nuclear background, with 95% confidence intervals. C. Clustermap of mean NARDINI pairwise blockiness Z-scores of eight amino acid residue classes for A′ UP, A′ DOWN and ITS2 proximal proteomes. D. Annotation analysis of RRM domain presence in A′ UP, A′ DOWN, and ITS2 proximal proteomes and nuclear control. Left: percentage of proteins containing at least one RRM domain. Right: percentage of proteins with a given number of RRM domains.

